# Machine learning-based classification of binary dynamic fluorescence signals reveals muscle stem cell fate transitions in response to pro-regenerative niche factors

**DOI:** 10.1101/2022.06.17.495556

**Authors:** Matteo Togninalli, Andrew T.V. Ho, Christopher M. Madl, Colin A. Holbrook, Yu Xin Wang, Klas E.G. Magnusson, Anna Kirillova, Andrew Chang, Helen M. Blau

**Affiliations:** Baxter Laboratory for Stem Cell Biology, Department of Microbiology and Immunology, Institute for Stem Cell Biology and Regenerative Medicine, Stanford School of Medicine, Stanford, California 94305-5175, USA; Department of Signal Processing, ACCESS Linnaeus Centre, KTH Royal Institute of Technology, 100 44 Stockholm, Sweden

## Abstract

The proper regulation of muscle stem cell (MuSC) fate by cues from the niche is essential for regeneration of skeletal muscle. How pro-regenerative niche factors control the dynamics of MuSC fate decisions remains unknown due to limitations of population-level endpoint assays. To address this knowledge gap, we developed a novel binary dynamic fluorescence time lapse imaging analysis (BDFA) approach that leverages machine learning classification strategies to track single cell fate decisions with high temporal resolution. Using two fluorescent reporters that read out maintenance of stemness and myogenic commitment, we constructed detailed lineage trees for individual MuSCs and their progeny, classifying each division event as symmetric self-renewing, asymmetric, or symmetric committed. Our analysis reveals that treatment with the lipid metabolite, prostaglandin E2 (PGE2), accelerates the rate of MuSC proliferation over time, while biasing division events toward symmetric self-renewal. In contrast, the IL6 family member, Oncostatin M (OSM) decreases the proliferation rate after the first generation, while blocking myogenic commitment. These insights into the dynamics of MuSC regulation by niche cues were uniquely enabled by our BDFA approach. We anticipate that similar binary live cell readouts derived from BDFA will markedly expand our understanding of how niche factors control tissue regeneration in real time.

## Introduction

Muscle stem cells (also known as “satellite” cells) play an essential role in skeletal muscle homeostasis, growth, repair, and regeneration during development and aging. Muscle stem cells (MuSCs) reside in an anatomically distinct compartment, between the myofiber plasma membrane and the basal lamina that surrounds the myofiber, which serves as an instructive microenvironment, or niche.^1,2^ MuSCs are of great interest as therapeutic targets due to their potent regenerative potential in repairing damaged tissue and restoring tissue function.^3–5^ The regenerative potential of MuSCs is due to their inherent ability to self-renew, proliferate, and differentiate to form myofibers that contribute to tissue repair. Quiescent MuSCs are defined by the expression of the paired-box transcription factor Pax7 and the absence of markers of “activation,” including cell cycle regulators and the Myf5 and MyoD transcription factors. Following activation, additional myogenic transcription factors, such as myogenin, govern further myogenic commitment and specialization. Upon activation, MuSCs expand through either asymmetric or symmetric division events, giving rise to self-renewing stem cells and to committed progenitors that undergo myogenic specification. Progeny of self-renewing division events can retain the capacity to return to a quiescent state upon resolution of muscle injuries, reconstituting the reserve pool of MuSCs.

During muscle tissue repair, MuSC fate is tightly controlled by networks of biochemical and biophysical cues. Dysregulation of these networks can lead to progressive loss of tissue function and to pathological conditions,^3^ as seen in the stem cell exhaustion that characterizes Duchenne muscular dystrophy.^6^ Our lab and others have identified a suite of soluble regulatory factors that is responsible for MuSC self-renewal, commitment to myogenic progression, or return to quiescence.^7–11^ In addition to soluble biochemical cues, insoluble biophysical cues from the MuSC niche, including extracellular matrix stiffness and composition,^12–18^ have been identified by our lab and others as crucial regulators of MuSC fate. Thus, both soluble biochemical cues and insoluble biophysical cues must be considered when assessing mechanisms regulating MuSC fate.

Here we leverage two key enabling technologies, bioengineered hydrogel culture substrates and single cell time lapse tracking,^12,13,19^ to explore the regulation of myogenic stem cell fate using a novel binary dynamic fluorescent reporter system and image analysis algorithm. This approach improves upon traditional experimental designs based on population-level analyses, which may obscure the properties of potentially interesting sub-populations and their growth dynamics in the averaged endpoint behaviors of global populations.^20^ To avoid these pitfalls, analyses tracking numerous single cells, which provide snapshots of entire cell populations at the single clone level, are desirable.^20,21^ However, while lineage tracking, molecular profiling and time-lapse imaging are attractive modalities of single-cell fate analysis,^22^ tracking single cell clones and their fate changes in real-time remains a challenge to-date, both due to the absence of accurate fluorescent reporter readouts and of algorithms for subsequent challenging data analyses.

Here we overcome prior limitations and enable monitoring of stem cell behavior and real-time cell fate transitions in culture, using a new strategy that permits spatial segregation and systematic manipulation of biomechanical and biochemical cues in conjunction with single cell lineage tracing in real time. Previously, we used retrospective analyses based on immunohistochemistry of cells fixed after time-lapse imaging is completed.^12,23^ Here, we provide a means of gaining insights into the dynamics of live stem and progenitor cell lineage analysis in conjunction with cell proliferation rate, survival, time to first division, time between divisions and cell migratory trajectories while maintaining genealogical relationships between daughter cells. We employed this binary dynamic fluorescence time lapse imaging analysis (BDFA) to evaluate the effects of PGE2 and OSM on MuSC fate determination kinetics using two fluorescent reporters that serve as proxies for stemness (Pax7 reporter) and commitment (myogenin reporter) and new machine learning algorithms, which allowed us to classify MuSC fate transitions. The combination of these signals captured by BDFA enabled the classification of MuSC fate into three distinct types: asymmetric, symmetric self-renewing and symmetric committed. BDFA revealed that the expansion of the stem cell pool in response to PGE2 occurs by increased numbers of symmetric self-renewing divisions across generations, while OSM elicits a quiescent-like phenotype by markedly reducing both the total number of divisions and the relative fraction of committed divisions, observations uniquely enabled by the ability to track cell fate decisions in single cells. We anticipate that our single cell BDFA approach will be broadly useful to map cell fate trajectories during development and regeneration of other tissue stem cell types.

## Results

### Population-level analysis of cell fate

We first established a population-level assay for cell fate decisions in response to treatment with pro-regenerative niche factors. For this purpose, we leveraged a myogenin reporter transgenic mouse model and synthetic hydrogels presenting biomimetic stiffness (*E* ∼ 12kPa) and laminin cell adhesive cues.^13^ Upon activation and initiation of myogenic commitment, characterized by expression of myogenin, the MuSCs harboring the reporter express Sun1-GFP. MuSCs from hind limb muscles were isolated by enzymatic digestion and FACS, using our established protocols.^12,24^ MuSCs were seeded onto hydrogel substrates and cultured for 7 days.

PGE2, a phospholipid metabolite generated from arachidonic acid at the onset of muscle injury, is a component of the body’s natural healing response. The niche factor PGE2 activates EP4 signaling in MuSCs to promote stem cell expansion essential to muscle regeneration.^7^ Because this niche factor is present during the initial wave of inflammation following injury, we added PGE2 to the cultured MuSCs at the time of seeding (**Fig. 1A**). Control cells were treated with vehicle (DMSO). A single early treatment with PGE2 was sufficient to increase cell number and bias the cells toward a more stem-like phenotype (**Fig. 1B**). By day 3 after treatment, an approximately 3-fold increase in cell number was observed (**Fig. 1C**), and by day 7, a significantly larger fraction of the MuSCs was positive for the muscle stem cell marker Pax7 and negative for the commitment marker myogenin compared to untreated control MuSCs (**Fig. 1D**). Thus, a population-level analysis of myogenic fate suggests that PGE2 treatment increases the number of cells retaining a stem cell phenotype.

**Figure 1.**
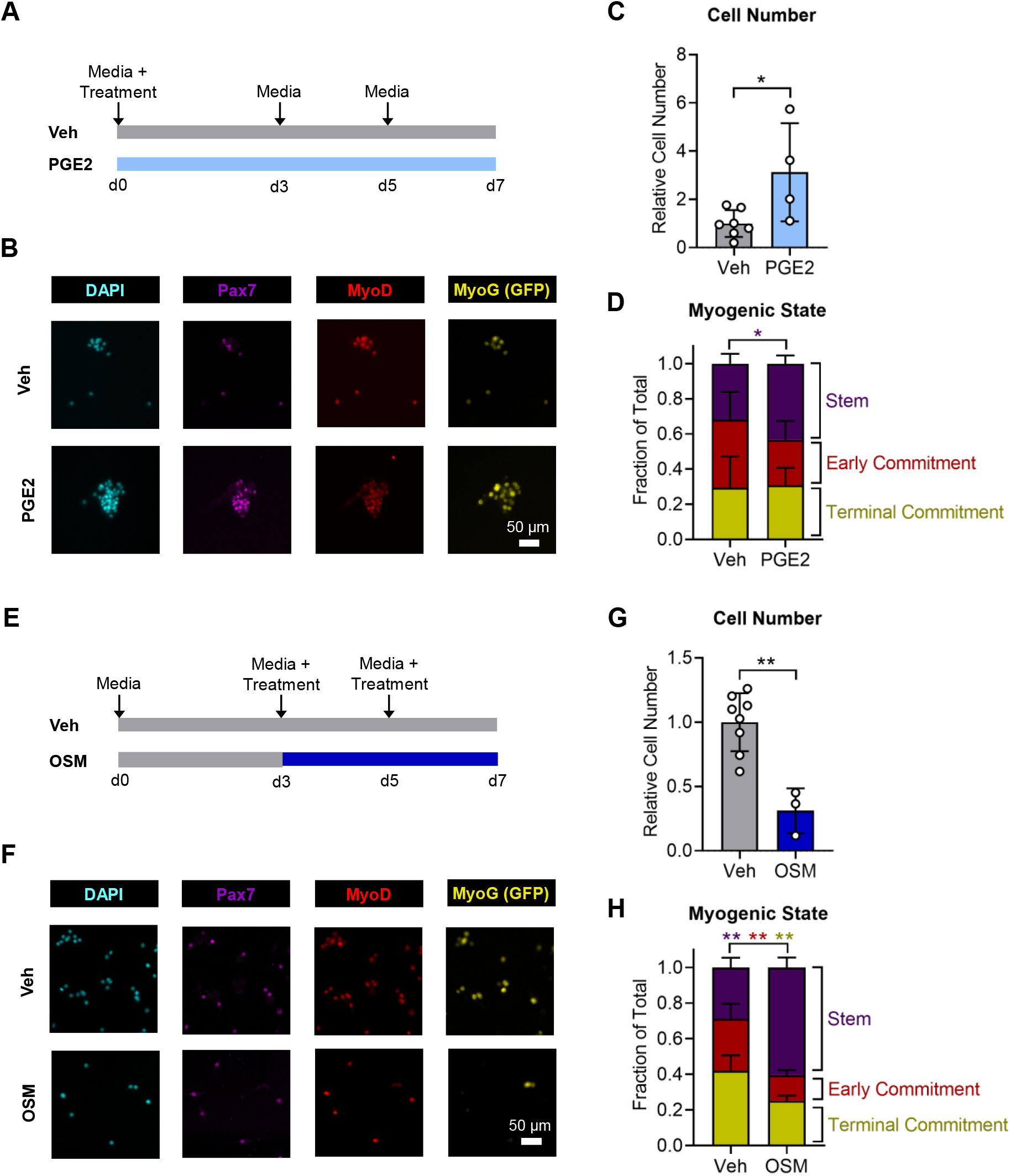
Population-level analysis of MuSC fate in response to pro-regenerative niche factors. **(A)** Schematic depicting the time course of MuSC treatment with PGE2. **(B)** Representative immunofluorescence images of MuSCs after 7 days of culture, treated with PGE2 or vehicle. **(C)** Quantification of cell number reveals increased proliferation in PGE2-treated MuSCs by day 3 in culture. **(D)** Quantification of myogenic factor expression 7 days after PGE2 treatment reveals expansion of the stem cell pool. **(E)** Schematic depicting the time course of MuSC treatment with OSM. **(F)** Representative immunofluorescence images of MuSCs after 7 days of culture, treated with OSM or vehicle. **(G)** Quantification of cell number reveals decreased proliferation in OSM-treated MuSCs at day 7 in culture. **(H)** Quantification of myogenic factor expression 7 days after OSM treatment reveals a bias toward stemness maintenance and decreased commitment in OSM-treated MuSCs. In D and H, cells that were Pax7+ and MyoG-were considered part of the “stem” population, cells that were MyoD+, Pax7-, and MyoG-were considered part of the “early commitment” population, and cells that were Pax7- and MyoG+ were considered part of the “terminal commitment” population. *p<0.05, **p<0.01, two-tailed Student’s *t*-test. Data in bar graphs are represented as mean ± standard deviation. In D and H, the color of the stars denoted the pairwise comparison performed (stem, early commitment, or terminal commitment). In C and D, n = 7 independent replicates for controls and 4 independent replicates for PGE2 treatment. In G and H, n = 8 independent replicates for controls and 3 independent replicates for OSM treatment.

OSM, a cytokine of the interleukin-6 family, plays a central role during muscle repair. It is secreted by macrophages, neutrophils and T cells and was shown to maintain MuSC fate and replenish the stem cell pool while regulating muscle regeneration.^8^ OSM is expressed in muscle during the later phases of regeneration, after MuSC activation.^8^ Accordingly, 3 days after seeding, MuSCs were treated with OSM or vehicle (PBS) control and cultured for an additional 4 days (**Fig. 1E**). Treated media was replenished at day 5 post-seeding. Treatment with OSM significantly suppressed the total number of cells by day 7 (**Fig. 1F,G**), with a significantly greater fraction of the OSM-treated cells staining positive for Pax7 and negative for the myogenin Sun1-GFP reporter relative to untreated control MuSCs (**Fig. 1H**). OSM treatment also reduced the fraction of cells in the early phase of myogenic commitment (positive for MyoD, but negative for Pax7 and myogenin) and in the terminal phase of myogenic commitment (positive for myogenin, but negative for Pax7) (**Fig. 1H**).

Therefore, based on population-level analysis, we postulated that PGE2 increases the number of self-renewing MuSCs whereas OSM acts as a pro-quiescence factor, reducing proliferation and myogenic commitment, while preserving stemness. However, resolution of the diverse effects of the two pro-regenerative regulators, PGE2 and OSM, on the dynamics of MuSC fate decisions was not possible by analyzing bulk populations.

### Binary fluorescent signals in single cell time lapse myogenic cell fate classification

Population-level analyses are limited by their reliance on averages taken at particular time points (**Fig. 1**), which obscure the contributions of subpopulations and the dynamics of cell fate changes.^20^ To overcome these limitations and gain further insights, we developed a new time lapse imaging approach to track fate changes in single MuSCs. We hypothesized that temporal analysis of binary signals derived from both the activity of a transcription factor highly expressed in stem cells (Pax7) and the activity of a reporter derived from the promoter of a transcription factor associated with myogenic commitment (myogenin) could yield sufficient information to allow the inference of division types and fate determination processes within the progeny of a single MuSC over time.

Four known transcription factors are commonly used to demarcate MuSC fate decisions and division kinetics (**Fig. 2A**).^25–30^ However, we reasoned that analysis of only two signals would suffice to delineate myogenesis into 5 distinct cell fates (**Fig. 2C**): (i) no division of a stem cell (Pax7-reporter^+^), (ii) asymmetric division to give rise to one Pax7-reporter^+^ stem cell and one MyoG-reporter^+^ cell, (iii) symmetric division giving rise to 2 Pax7-reporter^+^ stem cells, (iv) symmetric division giving rise to two MyoG-reporter^+^ cells, or (v) no division of a committed cell (MyoG-reporter^+^).

**Figure 2.**
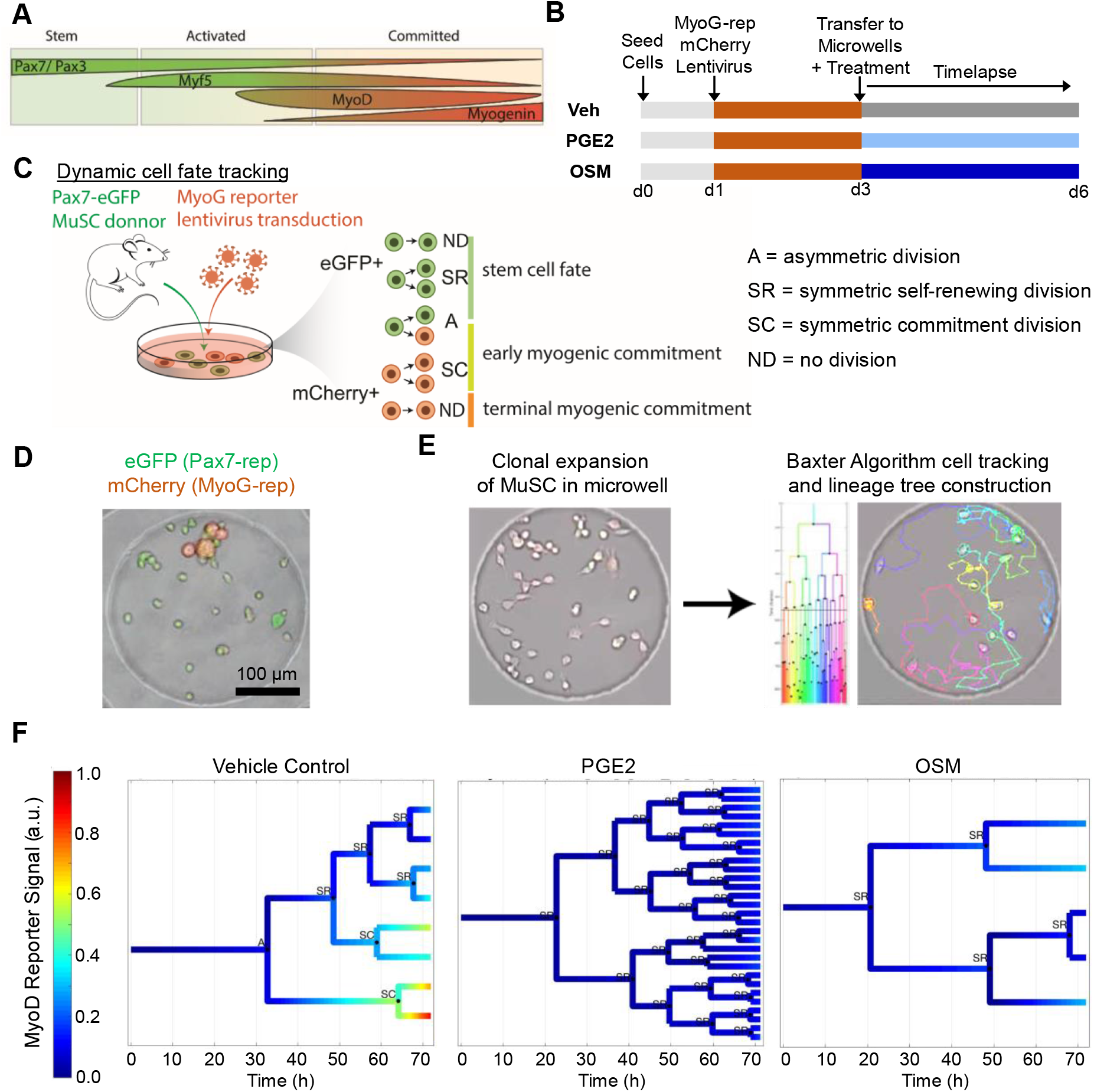
Development of a binary dynamic fluorescence time lapse imaging analysis (BDFA) approach to track MuSC fate decisions in real time. **(A)** Schematic depicting the relative timing of four transcription factors that determine MuSC fate. **(B)** Schematic depicting the workflow of binary reporter time lapse experiments to determine how PGE2 and OSM regulate MuSC fate decisions. **(C)** Graphic depicting the BDFA approach and how the binary fluorescent signals enable classification of cell fate decisions. **(D)** Representative fluorescence and brightfield micrograph of a single MuSC clone expressing the eGFP Pax7-reporter and the mCherry MyoG-reporter. **(E)** Single MuSCs seeded in hydrogel microwells were imaged over time and tracked using the Baxter Algorithm. **(F)** Representative lineage trees from vehicle, PGE2, or OSM treated MuSCs were generated by adding fluorescent reporter and machine learning classification modules to the Baxter Algorithm workflow. Division events were classified as symmetric self-renewing (“SR”), asymmetric (“A”), or symmetric committed (“SC”).

To measure the dynamics of fluorescence linked to transcription factor expression, we used MuSCs isolated from Pax7-eGFP mice,^31^ and infected the MuSCs in culture with a lentiviral-based reporter construct that expresses mCherry under the control of a genetic element derived from the proximal ∼1400 bp of the myogenin promoter (**Fig. 2B,C**; **Supplementary Fig. S1A-C**). We obtained a 59.33 ± 11.42% infection rate across 6 independent replicate experiments (**Supplementary Fig. S1D**). MyoG reporter activity was validated by treating the transduced cells with differentiation media to induce myogenic commitment and measuring mCherry fluorescence by microscopy. As expected, differentiation media induces mCherry expression in transduced cells but not in sham control cells (**Supplementary Fig. S1E**). Furthermore, no detectable virus-induced cytotoxicity was observed by staining for the apoptotic marker cleaved caspase-3 (**Supplementary Fig. S1F**).

The resulting MuSCs with dual fluorescent markers were then seeded at low density onto the 300 μm diameter microwells formed in 12 kPa hydrogels to obtain on average one single MuSC in each microwell.^12^ A MuSC clone that expressed both Pax7-GFP reporter and MyoG-mCherry reporter post-infection in a hydrogel microwell was readily visualized by fluorescent time lapse microscopy (**Fig. 2D**). The initial positions of each clone were manually selected under the time lapse microscope to ensure the first frame of the image sequence captured a single Pax7-GFP^+^ MuSC. We acquired bright field images at 5-minute intervals and fluorescence images of the GFP and mCherry probes at 20-minute intervals to reduce phototoxicity while maintaining sufficiently high temporal resolution. To ensure that the tracked clones also harbor the myogenin reporter construct, we differentiated the MuSC clones at the end of the imaging timepoint and confirmed the presence of mCherry fluorescent reporter. Clones that did not harbor cells that expressed myogenin were excluded from the analysis.

### Methods developed to augment automated cell tracking and fate classification

While several examples of automated cell tracking software have been reported,^19,32,33^ none has provided a solution that allows the identification of cell fate in real-time. To overcome this challenge, we built upon our previously described and well-validated Baxter Algorithm.^19,34^ We augmented the existing Baxter Algorithm by adding fluorescence analysis and cell division type classifier scripts used for the automated tracking of a single MuSC colony. As validation, we performed test runs with manual correction on every tracked well to train the algorithm, an essential parameter in the workflow sequence to quantify the fluorescence signal generated by cells. The resulting Baxter Algorithm allowed for complete lineage tree reconstruction coupled with fluorescent reporter expression dynamics and division type classification (**Fig. 2E**).

We further increased the performance of the time-lapse acquisition system by including fluorescence image homogenization steps to account for the uneven illumination conditions, which yielded comparable imaging data across samples (see **Methods**). Once the image sequence was homogenized, the average fluorescence pixel value was extracted from within the cell borders generated by the tracking algorithm and further normalized by applying a moving average filter. Finally, in order to generate comparable data across colonies and experiments, the quantified fluorescence signals of the Pax7 and MyoG reporters were normalized with respect to the maximum and minimum values observed in each microwell during the imaging period and after differentiation (**Supplementary Fig. S2A**). Since the raw fluorescent images acquired from time lapse microscopy exhibited a continuum in fluorescent signals, distinct cell fates could not be classified based solely on the numerical values of the fluorescent signals. Therefore, the signals were subsequently classified into 5 quintiles to allow visual separation of distinct phases and still maintain the characteristics of the expression of each tracked clone (**Supplementary Fig. S2A**).

To enable classification for the type of cell division that had occurred using the processed fluorescent images, we generated an additional script that enables analysis of the binary fluorescent signals from the time lapse image sequences. This analysis yielded the automated generation of a lineage tree with a heatmap overlay representing the expression of the MyoG reporter over time (**Fig. 2F**). Furthermore, this appended script also generated the division types for each cell division event without user input by implementing the 1-nearest neighbor (1-NN) classifier applied to more than 220 k-means-derived and manually-validated training samples (**Supplementary Fig. S2B**). Thus, each division was classified into one of the following scenarios for individual MuSCs: (i) no division (Pax7-GFP^+^), (ii) asymmetric division (designated as A) to give rise to one Pax7-GFP^+^ stem cell and one MyoG-mCherry^+^ cell, (iii) symmetric division giving rise to 2 Pax7-GFP^+^ stem cells (designated as SR for “symmetric renewal”), (iv) symmetric division giving rise to two MyoG-mCherry^+^ cells (designated as SC for “symmetric commitment”), or (v) death (**Fig. 2C,F**). To benefit fully from the large quantity of information generated by the time-lapse analysis and to consider all the acquired information related to cell division, we included 38 additional identifiable features for every division as input parameters to refine our cell fate classifier. These parameters include features such as parent/daughter cell minimum/maximum/average/variance of Pax7 and MyoG reporter expression as well as the lifetime difference of two daughter cells, i.e., the time each of the two cells existed, and their ability to further divide (**Supplementary Table S1**).

Since assigning division-type is a classification problem, we employed a machine learning approach.^35^ A K-means clustering algorithm (k = 3 for symmetric self-renewing with two stem daughter cells, symmetric committed with two committed daughter cells, and asymmetric divisions) coupled to a 1-nearest neighbor algorithm were used to create three classes corresponding to the three division types and to assign new divisions to these classes. A few divisions (n=12 from all experimental conditions) with an obvious division type were manually annotated and used as ground-truth values for assignment of weights of the features during clustering (**Supplementary Fig. S2B**). The resulting augmented Baxter Algorithm generated full lineage tree representation, including MyoG (or Pax7) reporter expression and division type, providing insights correlating cell fate, cell division and cell behavior in a manner that cannot be achieved by bright field time-lapse imaging analysis. Furthermore, the software developed allowed us to acquire the fluorescence expression values for each generation of cell progeny starting from the stem cell, enabling a more precise evaluation of transcription factor dynamics at the single clone level.

Taken together, the automated tracking of the fate of a single MuSC colony presented here benefitted from the inclusion of fluorescent stem cell and fluorescent committed stem cell reporter expression analysis and cell division type classifiers analyzed by newly developed machine learning algorithms. These new features enabled dynamic analysis of lineage tree reconstruction in a manner previously not possible.

### Elucidating the role of PGE2 and OSM in regulating MuSC fate dynamics

We sought to determine how the pro-regenerative niche factors PGE2 and OSM influence clonal proliferation and fate dynamics in MuSCs. We previously showed that both PGE2 and OSM increase the engraftment efficiency of transplanted cells and enhance stem cell-mediated regeneration of tissue damage.^7,8^ However, the mechanisms at the level of cell fate determination were not fully resolved in these studies. To achieve this, here we used the BDFA pipeline to automate cell tracking and fate classification. We used Pax7-eGFP reporter MuSCs isolated by FACS and transduced them with the lentiviral MyoG-mCherry reporter construct. In contrast to the endpoint population-level assays described above, the combination of two live fluorescent reporters enabled determination of MuSC fate decision dynamics on a single cell level. The transduced cells were seeded on microwell hydrogels and treated with vehicle (control), PGE2, or OSM for time lapse imaging. Brightfield and fluorescence images were collected over the following 72 hours, as described above. After completion of the time lapse imaging, the expression of the MyoG-mCherry reporter construct was validated by inducing differentiation in the tracked cells. Clones harboring the MyoG reporter were then analyzed by the optimized BDFA algorithm.

Our optimized analysis pipeline generated lineage trees for each condition that were overlayed with the relative MyoG reporter signal and cell division classifications (**Fig. 2F**). Two striking patterns emerged. In PGE2 treated cells, the number of division events during image acquisition was greater than controls, and the division events were skewed toward symmetric self-renewal compared to controls. In OSM treated cells, the total number of divisions was decreased, but comparatively more of the divisions were symmetric self-renewing versus controls. A major advantage of the time lapse approach is the ability to track single cells over time. Our analysis tracking the number of live and dead cells in each generation (**Fig. 3**) reveals a rapid progression through successive generations in the control and PGE2 treated cells, indicating consistent proliferation that is more rapid in the PGE2 treated condition. In contrast, the OSM treated cells spend longer in each generation and do not divide as frequently, consistent with the reported role of OSM as a pro-quiescence factor. All conditions exhibited relatively little cell death.

**Figure 3.**
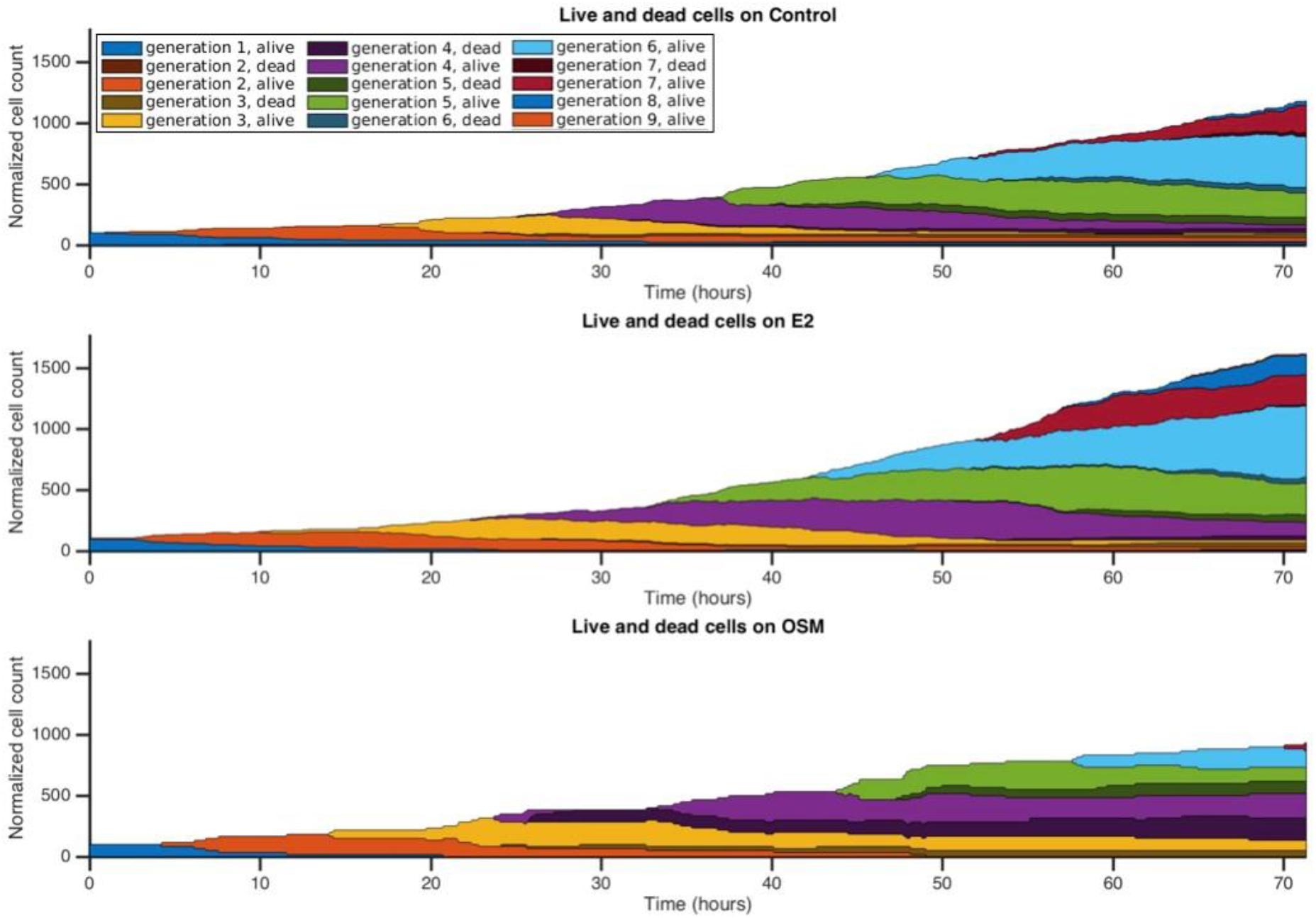
Tracking single MuSCs across generations using time lapse imaging. The BDFA approach enables tracking of individual MuSCs and their progeny through numerous rounds of cell division. Treatment with PGE2 accelerates cell division, with some microwells having progeny in the 9^th^ generation after 3 days. In contrast, OSM slows down proliferation, acting as a pro-quiescence factor. At least 18 independent clones were tracked for each condition from at least 2 independent experimental replicates.

The division-type classification associated with each division event enabled facile assessment of the cell fate dynamics in response to PGE2 and OSM treatment over time. PGE2 treatment results in greater overall proliferation than control, while OSM results in decreased proliferation (**Fig. 4A**). Importantly, our temporally resolved single cell data enable us to observe how proliferation rate changes over time, with PGE2 treated cells showing consistent or increased proliferation rates, while OSM treated cells exhibit decreased proliferation rates. However, when considering the myogenic fate of the proliferating cells, both PGE2 and OSM enhance the relative fraction of divisions classified as symmetric self-renewal. Thus, both PGE2 and OSM bias the cells toward a pro-stem cell phenotype while attenuating myogenic commitment. The generational breakdown of division types emphasizes the constant distribution of symmetric division types in the PGE2 culture, wherein a ratio of approximately 60% self-renewing, 15% asymmetric and 25% committed divisions is maintained throughout the first four generations, resulting in a steady increase in MuSC proliferation (**Fig. 4B**). On the other hand, the OSM treated cells showed a comparable self-renewing division distribution to controls only in the first generation, and cell division rate declined with subsequent generations. Thus, our BDFA approach revealed that PGE2 treatment expands the stem cell pool by increasing the rate of symmetric self-renewing divisions, while OSM treatment drives a quiescent-like phenotype by decreasing the rate of divisions that are biased toward symmetric self-renewal.

**Figure 4.**
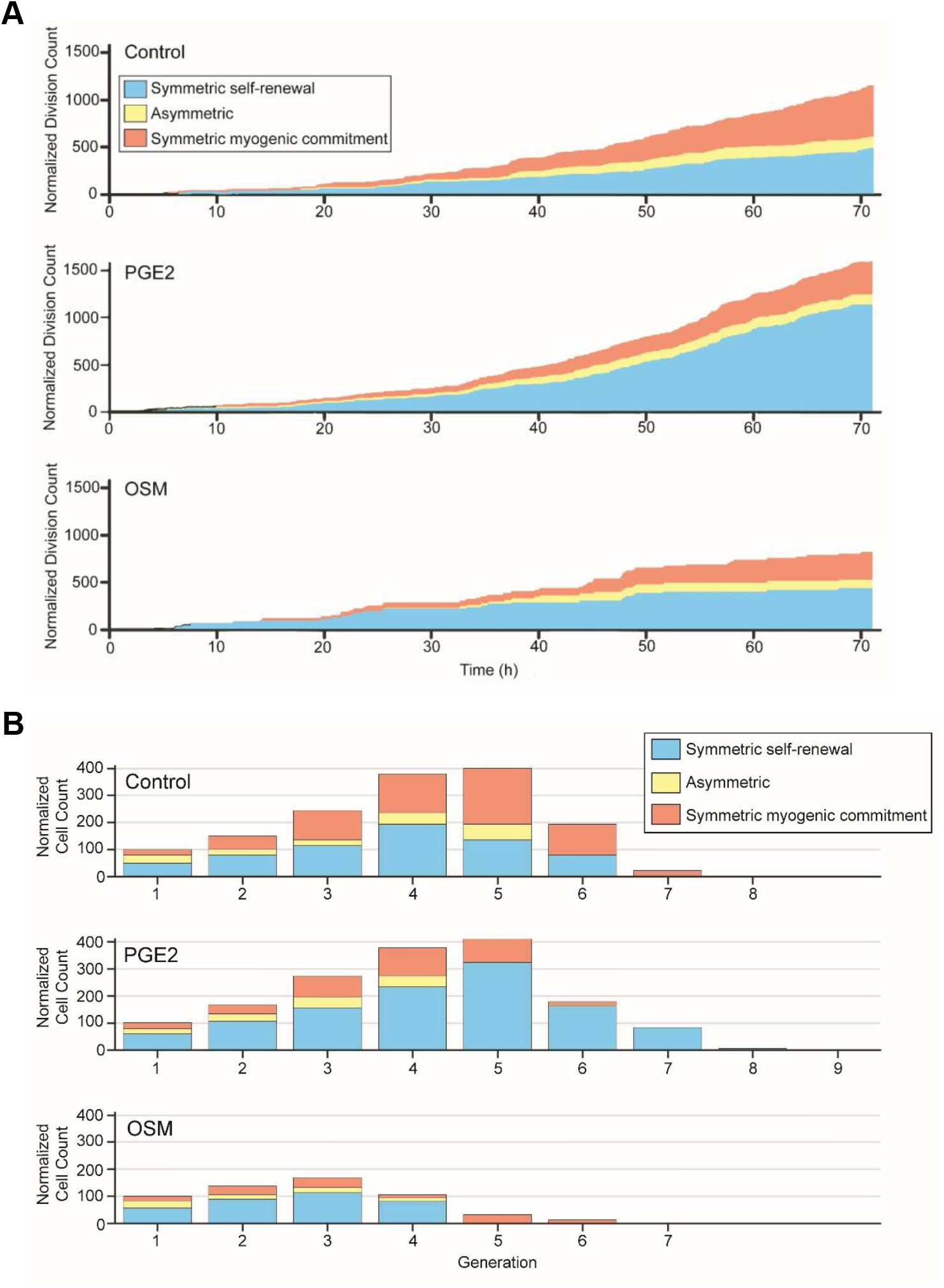
Machine learning-based division classification reveals cell fate dynamics. **(A)** Cumulative division counts and classifications for vehicle, PGE2, and OSM treated MuSCs, normalized to represent the same number of starting clones per condition. **(B)** Classification of division types broken down by generation reveals temporal dynamics of cell fate decisions. To enable comparisons across conditions, the number of initial clones was normalized to be 100 in each condition. At least 18 independent clones were tracked for each condition from at least 2 independent experimental replicates.

## Discussion

Muscle repair following injury is orchestrated by networks of niche factors that direct muscle stem cell self-renewal, activation, expansion, and differentiation.^2,26,27^ Our understanding of how these factors impact individual myogenic cell fate decisions in MuSCs is frequently limited by the use of population-level assays taken at specific time points during regeneration in vivo or following MuSC isolation and culture in vitro. To overcome the limitations of population-level fixed time point assays, we developed a method for quantification and analysis of binary fluorescence signals during time lapse imaging of muscle stem cells and applied it to an automated system for classifying identified cell divisions via machine learning strategies. By leveraging K-means clustering approaches applied to 38 distinct parameters derived from our time lapse data and a 1-nearest neighbor classifier, we were able to robustly classify cell fate decisions without tedious manual and subjective scoring by blinded observers. This approach enabled a full reconstruction and comparison of muscle stem cell fate determination processes.

Our approach using live cell time lapse imaging offers substantial benefits as a non-invasive method for elucidating the dynamics of cell fate determination. Unlike end point approaches, time lapse imaging does not entail the destruction of cells upon measurement. It allows for lineage tree reconstruction, dynamic evaluation of dynamic molecular changes via fluorescent reporters, and identification of cellular heterogeneity. Furthermore, our BDFA approach yields markedly increased temporal resolution, since it does not rely on time-interval snapshots like other types of single-cell analysis previously used by us and others. Fluorescence time lapse imaging is currently the optimal technique for quantifying dynamic molecular changes in live cells and evaluating short-term single stem cell fate determination in vitro. In conjunction with our machine learning-aided pipeline, fluorescence time lapse imaging maximizes the utility of time lapse data by automating cell fate classification.

We applied our BDFA approach to investigate how two pro-regenerative niche factors, PGE2 and OSM, regulate MuSC fate determination. Cell tracking revealed changes in proliferation rates, while fluorescent reporters and cell division characteristics provided novel insights into the dynamics of the fate determination process. Treatment with PGE2 accelerated the rate of symmetric self-renewing divisions, expanding the stem cell pool, providing insights into the mechanism by which PGE2 boosts MuSC-mediated muscle repair.^7^ Treatment with OSM slowed the overall cell division rate, while biasing those divisions that did occur toward symmetric self-renewal, consistent with its role as a pro-quiescence factor, as previously described.^8^ Identification of these temporal changes in proliferation kinetics and cell fate decisions would not be possible without our single cell BDFA approach. Dynamic fluorescence time-lapse is therefore an advantageous new tool for stem cell fate elucidation and a resource for drug screening assays. We anticipate that further application of BDFA to other cell types and pro-regenerative factors will enable previously inaccessible insight into the dynamics of fate decisions underlying stem cell-mediated regeneration.

## Experimental Procedures

### MuSC isolation

#### Cell extraction

All experiments and protocols were performed in compliance with the institutional guidelines of Stanford University and Administrative Panel on Laboratory Animal Care (APLAC). Cells were obtained from either MyoG-CreERT2 Sun1-GFP or Pax7-eGFP mice bread as previously described and aged between 2 and 7 months.^13,31^ The hindlimb muscles were removed from both legs of each mouse, washed in Phosphate-Buffered Saline (PBS) and placed in C-tubes (gentleMACS C Tubes, Miltenyi Biotec Inc., San Diego, CA) and digested with collagenase and dispase as previously described.^12,24^ The resulting mixture was filtered through a 45 µm cell filter (Thermo Fisher Scientific, Waltham, MA).

#### Fluorescence Activated Cell Sorting (FACS)

The MyoG-CreERT2 Sun1-GFP MuSCs were stained with PE-Cy7 conjugated antibodies against CD45, CD11b, CD31, and Sca1 (BD Biosciences; lineage markers, Lin) to label non-muscle cell types, and fluorescently tagged antibodies to label MuSCs: integrin-α7-PE (Ablab) and CD34-Alexa647 (BD Biosciences). The cell suspension was sorted by FACS to deplete non-muscle lineage cells and enrich MuSCs (Lin-/integrin-α7+/CD34+/ GFP-). The Pax7-eGFP reporter cells were sorted by gating on the GFP signal (repeatedly obtaining a purity > 88%) and plated in collagen-coated well plates (Corning Incorporated, Corning, NY).

#### Cell culture

For population-level analyses, MuSCs were cultured in Fluorobrite DMEM with 15% fetal bovine serum (FBS), 1% antibiotic/antimycotic, 1% non-essential amino acids, 1% L-glutamine, 1x sodium pyruvate (Thermo Fisher Scientific), and 2.5 ng ml^-1^ basic fibroblast growth factor (bFGF). For lentiviral transduction and reporter construct optimization, MuSCs were cultured in 45% low glucose DMEM, 40% F10 nutrient mix (Thermo Fisher Scientific), 15% fetal bovine serum (FBS), 1% antibiotic/antimycotic, and 2.5 ng ml^-1^ bFGF. Medium with the same composition but using FluoroBrite DMEM was used for time-lapse imaging. MuSCs were maintained at 37°C in 5% CO_2_ and medium was changed as noted in figures. To induce differentiation, medium containing 2% horse serum and 1% antibiotic/antimycotic in FluoroBrite DMEM was added.

### Viral infection

#### Viral vector construct

To observe myogenic commitment, the cells were infected with a viral construct containing mCherry driven by a ∼1400 bp region of the myogenin promoter. The viral vector was based on a commercially-available pEZX-LvPM02 lentiviral construct containing the myogenin promoter (MPRM15676-LvPM02, Genecopoeia, Rockville, MD). The plasmid map can be found in **Supplementary Fig. 1C**.

#### Transduction

Viral transduction occurred the day after MuSCs were plated. Polybrene (TR-1003-G, EMD Millipore, Hayward, CA) was added to the wells to increase the efficiency of viral transduction. A concentration of 40 µl per 1 ml of medium was used. The medium was changed 4 hours after infection to increase cell viability. After transduction, the cells were cultured for 48 hours before being seeded onto the hydrogel microwells.

#### Cell viability

Cell viability was measured 2 days after transduction in cells with and without virus by staining for cleaved caspase-3 as described below.

#### Infection rate

Infection rate was calculated as follow

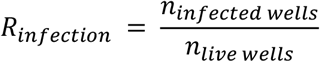

Where *n_infected wells_* and *n_live wells_* are the number of wells in which the initial clone generated a colony expressing mCherry after differentiation (day 5) and the number of wells in which the colony survived respectively. The dead colonies were not considered for this analysis because such colonies most often result from apoptosis of the initial MuSC, and the MyoG reporter is not expressed at that early time point, making it impossible to distinguish between false and real negatives.

### Hydrogel preparation for population-level assays

Uniform hydrogels with elastic moduli of ∼12 kPa were prepared in 24-well glass bottom tissue culture plates as previously described.^13^ Briefly, 8-arm 10 kDa PEG-azide was mixed with a solution of 4-arm 10 kDa PEG functionalized with bicyclo[6.1.0]nonyne (BCN) and purified laminin functionalized with 3.5 kDa linear PEG azide at a final polymer concentration of ∼5.4% w/v. The gel precursor solution (200 µL) was added to each well, and the plate was centrifuged at 2,000 g for 15 minutes at room temperature to afford a thin layer of hydrogel covering the bottom of the well. Gels were further incubated at 37°C for 30 minutes to complete crosslinking, washed with PBS plus 1% antibiotic/antimycotic, and equilibrated in cell culture media at 37°C prior to cell seeding.

### Hydrogel microwell preparation

#### Hydrogel precursor preparation

Poly(ethylene glycol) (PEG) hydrogels were designed to have an elastic modulus of ∼12 kPa.^12^ The hydrogels were prepared from two multi-arm precursors: 4-arm 10kDa PEG-thiol (PEG-SH; Sunbright PTE-100SH, NOF Corporation, Tokyo, Japan) and 8-arm 10 kDa PEG-vinyl sulfone (PEG-VS; Sunbright HGEO-100VS, NOF Corporation, Tokyo, Japan). PEG-VS was dissolved in sterile-filtered 0.2M triethanolamine (TEOA, pH 8) to 10% w/v and filtered through a 0.45 µm syringe filter. PEG-SH was dissolved in sterile-filtered distilled water at a concentration of 10% w/v solution and filtered through a 0.45 µm syringe filter. Stock solutions of precursors were stored at −20°C.

#### PDMS stamp preparation

To generate microwell structures in the hydrogels, poly(dimethylsiloxane) (PDMS) stamps were fabricated by casting PDMS precursors (Sylgard 184 Silicone Elastomer, Dow Corning Corp., Midland, MI) on silicon wafers with 300 µm diameter microwells, generated via photolithography, and cut to fit regular 24-wells plates.^12^ The resulting PDMS stamps had cylindrical micropillar structures. To functionalize the hydrogels microwell bottoms, the tops of the cylindrical micropillars on the stamps were coated with laminin (Roche, ref 11243217001). The laminin was dialyzed in phosphate-buffered saline (PBS) and deposited on Bis-Tris gels (NuPAGE, Thermo Fischer Scientific) previously washed in PBS and dried. The PDMS stamps were then deposited with the feature side down onto the dried gels for 30 minutes to transfer the laminin to the PDMS.

#### Hydrogel fabrication

The hydrogel precursors were degassed and kept on ice. Triethanolamine (TEOA) was used as a buffer for the crosslinking reaction. PEG-SH and PEG-VS were sequentially added to the TEOA to reach a 3.4% wt/vol gel concentration and were vortexed. The laminin-coated PDMS stamps were deposited on glass slides with the features facing up, 200 µl of hydrogel mixture was added to each stamp, and a second glass slide was placed on top of the PEG solution. Spacers with a fixed height located on each side of the PDMS stamps and hydrogel mixture ensured homogeneous thickness of the formed gels. The crosslinking hydrogels were then incubated for 18 hours at 37 °C. The hydrogels were then removed from the slides and PDMS and hydrated in PBS for 8 hours at 4°C, washed and stored in PBS at 4°C.

#### Cell seeding

Hydrogels were affixed to the bottom of 24-well plates with a mixture of gel precursors (3.4% w/v, as above). The gels were washed with PBS and growth medium and allowed to equilibrate in growth medium for one hour. The cells were detached from their plate with Trypsin-EDTA 0.5% (15400054, Thermo Fisher Scientific) and diluted to appropriate concentrations (1000 cells/ml for ø 300 µm microwells) prior to seeding. These concentrations ensure that a sufficient number of wells contain a single cell.^12^ E2 (10 ng/mL, Cayman Chemical, Ann Arbor, MI) or OSM (100 ng/mL, R&D Systems, Minneapolis, MN) was then administrated, and time lapse imaging was started. Every replicate consisted of one treated and one control well.

### Time lapse setup

The plates containing the single cells in hydrogel microwells were positioned in the environmental chamber of an inverted microscope with motorized stage and temperature-controlled chamber (Axio Observer Z1 with PM S1 incubator, Carl Zeiss AG, Oberkochen, Germany). The chamber was equipped with a gas exchanger (CO_2_ module S, Carl Zeiss AG) and was kept at 37.5°C and 5% CO_2_. The images were acquired with an AxioCam MRm camera (Carl Zeiss AG) at 1388×1040 (brightfield) or 4×4 binning 346×260 (fluorescent channels) pixel resolution. The microscope software AxioVision (Carl Zeiss AG) was set up to acquire bright field images in different positions at an interval of 5 minutes and a custom-made script was written to acquire fluorescence images of the same positions at an interval of 20 minutes using Filter Set 44 (Carl Zeiss AG) for GFP and Filter Set 20 (Carl Zeiss AG) for mCherry. The acquisition was conducted for a period of 3 days (72h). The exposure times for the different channels (15-50 ms for brightfield, 80-160 ms for GFP and 100-200 ms for mCherry) and the camera parameters were carefully chosen to reduce phototoxicity. Each single-cell colony was manually selected, and its position was recored by the software before starting the time lapse, with an equal number of wells imaged for each condition (drug treatment and control). Each time-lapse experiment produced about 45 image sequences in total. The resulting images were analyzed with an augmented version of the Baxter Algorithm in Matlab (MathWorks, Natick, MA).^19^ After the time lapse acquisition, differentiation medium was added to the well plate to induce myogenic commitment to validate that the tracked clones were successfully transduced with the MyoG reporter. Validation images of differentiating cells were taken 2 days (48h) after addition of differentiation medium. Only clones expressing mCherry after the differentiation step were further analyzed.

### Time lapse algorithm

#### Existing algorithm

The existing algorithm used in the Baxter Laboratory is a batch algorithm that uses all available images to perform individual track linking by using temporal information in a probabilistic manner. The current algorithm separates the image analysis into two parts: image segmentation and track linking. It is able to handle false positive detections (small non-cellular objects), cell clusters and missed detections. The algorithm successfully keeps track of cell movement, division, and cell death, as well as cells entering and exiting the image window. To do so, the algorithm starts with the hypothesis that no cells are present on the image sequence. It then adds one optimal track at a time to maximize a probabilistically motivated scoring function. The added track’s optimality is guaranteed by solving a dynamic programming problem using the Viterbi algorithm. The algorithm stops once the scoring function cannot be further maximized. The cell tracking component relies entirely on the bright field images, due to the higher acquisition frequency for higher tracking precision. The computer-generated tracks of every selected well (MyoG reporter positive after differentiation) were corrected by hand as needed to ensure the most reliable quantified data.

#### Improvements

The existing version of the algorithm was augmented with a fluorescence data analysis plug-in. The Baxter Algorithm was updated to handle fluorescence channels and to integrate data derived from these channels into the final analysis. The new plug-in allows for extraction of fluorescence data from the cell regions identified by the tracking algorithm, as described in more detail below.

### Fluorescence analysis

#### Image retrieval

A Python-based script was written to rename and resize all the binned images obtained from the acquisition so they could match the existing requirements of the Baxter Algorithm.

#### Signal treatment

In order to counter uneven fluorescence illumination present in each image sequence, pre-processing was applied to all fluorescence images and their respective extracted signal. An initial homogenization step was applied to the image sequence to limit large fluctuations of signal between time-adjacent images (flickering): the average signal of every fluorescence image was computed and used to fit the sequence average intensity throughout time. The average of every image was then brought to their computed value by single-value subtraction (or addition). A power law was chosen to fit the average signal of the GFP fluorescent channel due to the residual medium auto-fluorescence caused by ambient illumination during the time lapse setup, whereas a linear polynomial fit was used for the mCherry signal, since auto-fluorescence was not observed in that channel. Image homogenization was preferred over simple background subtraction to account for the varying nature of the fluorescence image averages. Furthermore, adding a single value to every pixel was preferred over multiplying them by a constant in order not to excessively amplify noise.

#### Fluorescence signal

The average signal for every cell was then extracted from their respective cell border (obtained from the tracking algorithm) in every image of the image sequence and further smoothed with a moving average filter (with window size of 5). The few overexposed images taken on the mCherry channel, due to the delay of the microscope shutter, were discarded as source of fluorescence signal. Finally, the signal was normalized with respect to the minimum and the maximum signal of the image sequence to have comparable data among colonies and across experiments. The maximum value was taken as either the 99 percentile of the differentiated cells image taken after two days in differentiation medium or the maximum value observed during the time-lapse, whichever was higher.

#### Differentiation visualization

The normalized fluorescence signal for a cell was classified in five quintiles for representation purposes. This representation was chosen since cells tend to have continuous levels of expression for the two signals and do not follow a simple on/off type of behavior.

### Division classification

#### Rationale

The task of classifying observed divisions is central to the investigation of dynamic cell fate determination. A first-pass approach consisted of hard-coding the conditions a cell had to match in terms of quantified MyoG reporter pattern, but predicting all possible behaviors is complex. Furthermore, time lapse analysis yields a large quantity of other data in which division type-related information could be hidden. It would be overly simplistic not to consider the lifetime difference of two daughter cells, or their ability to further divide. To harness the value of all available information, machine learning techniques were applied to the collected data.^35^

#### Features extraction

The previously described algorithm generated objects containing all relevant data for every cell. Building on top of that, features corresponding to each division (i.e., reflecting the parent and daughter cells behaviors) were extracted and normalized so as to have similar variances and values across features. Weights were then assigned to each feature to match their importance in characterizing a cellular division. Weights were decided empirically and validated with ground truth lineage trees. In total, 38 features were collected from every division. A complete list can be found in **Supplementary Table 1**.

#### K-means clustering

Divisions of cells were classified using the k-means clustering algorithm in Matlab® combined with the 1-nearest neighbor classifier. K-means clustering is a method used in unsupervised or semi-supervised learning to partition *n* observations in *k* clusters. The method finds *k* centroids so as to minimize the distance of every observation to its cluster centroid. It allows regrouping of similar observations together without the need for labeled samples (i.e., samples with explicit, manually-assigned ground truth). According to the elbow method – commonly used to identify the best suited number of clusters for a dataset – 4 clusters would represent the overall data better than 3, but the only further distinction of a fourth class would be to indicate new divisions that do not yield dividing cells at the end of a time-lapse (where it is not possible to know whether cells will undergo further divisions). The clusters were therefore obtained on a specific set of sequences (N = 12 sequences, n = 229 total divisions).

#### Division classification

All new divisions that were not used in the initial clustering were then classified using a 1-nearest neighbor algorithm that assigned every division to its closest cluster centroid. The features for these new divisions underwent the same normalization steps that the features of the division used in the clustering part. It is important to note that this classification is not perfectly accurate for borderline cases but generates comparable results across clones.

#### User interface

All the above-mentioned image processing and data analysis steps were integrated in a new analysis window under the Baxter Algorithm software. Seamless integration of the different software components was ensured in Matlab®. New settings were added to the list of settings in the Baxter Algorithms to accommodate parameters such as mismatching intervals between brightfield and fluorescent image acquisition and measurements after differentiation. The new user interface allows for direct output of statistics about single cell lineage tree or overall population statistics.

### Videos

Videos were compiled in Photoshop (Adobe, San Jose, CA) using the frames generated by the algorithm. The fluorescence signal was enhanced for visualization and quantification (**Supplementary Videos 1-3**).

### Immunostaining

To assess viral toxicity, cells were fixed with 4% paraformaldehyde for 10 minutes and washed in PBS. A blocking solution of 1% bovine serum albumin (BSA) and 0.05% Triton X-100 in PBS was applied, and then cells were stained with rabbit cleaved caspase-3 primary antibody (Santa Cruz Biotechnology, sc-7148, 1:200) and Alexa Fluor 647-conjugated goat anti-rabbit IgG secondary antibody (Jackson Immunoresearch Laboratories, West Grove, PA, 1:500) after viral transfection to assess any potential viral toxicity. For population-level analysis of cell fate, cells were fixed with 4% paraformaldehyde in PBS for 30 minutes, washed three times with PBS, permeabilized with PBS plus 0.25% (v/v) Triton X-100 (PBST) and blocked with 5% bovine serum albumin (BSA) and 5% goat serum (GS) in PBST. Cells were subsequently incubated with primary antibodies against Pax7 (Developmental Studies Hybridoma Bank, 2 μg/mL) and MyoD (Santa Cruz Biotechnology, clone G-1, 1:200) diluted in 2.5% BSA and 2.5% GS in PBST. Samples were washed three times with PBST and then incubated with secondary antibodies (Cy3 goat anti-mouse IgG2b and AF647 goat anti-mouse IgG1, Jackson ImmunoResearch, 1:500) diluted in 2.5% BSA and 2.5% GS in PBST. In all cases, nuclei were counter-stained with Hoechst 33342 (Invitrogen, Thermo Fischer Scientific, 1:1000). Images were acquired on an Axio Observer Z1 inverted fluorescence microscope.

## Acknowledgments

The authors would like to thank Dr. Matthias Lutolf for helpful discussions about the BDFA approach. M.T. was supported by a Zeno Karl Schindler grant and a Swiss Study Foundation Scholarship. A.T.V.H. was supported by a Muscular Dystrophy Association (MDA) Career Development Award (MDA 217821). C.M.M was supported by a Life Sciences Research Foundation Postdoctoral Fellowship and a U.S. National Institutes of Health (NIH) K99 award (K99 AG071738). Y.X.W. was supported by the Canadian Institutes of Health Research (MFE-152457) and an NIH K99 award (K99 NS120278). This study was supported by funding from the Baxter Foundation, the Li Ka Shing Foundation, and the NIH (R01 AG069858 and R01 AG075436 to H.M.B.).

## Author Contributions

M.T., A.T.V.H., C.M.M., and H.M.B. conceived of the study, designed experiments, and wrote the manuscript. M.T., A.T.V.H., C.M.M., C.A.H., and Y.X.W. performed experiments. M.T., A.T.V.H., C.M.M., C.A.H., Y.X.W., K.E.G.M., A.K., and A.C. analyzed data. H.M.B. is the principal investigator.

## Competing Interests

The authors declare the following financial interests/personal relationships which may be considered as potential competing interests: A.T.V.H. and H.M.B. are named inventors on patent applications held by Stanford University regarding PGE2 and muscle regeneration licensed to Myoforte Therapeutics. H.M.B. is a cofounder of Myoforte Therapeutics, and receives consulting fees and has equity and stock options from Myoforte Therapeutics. H.M.B. is a cofounder of Rejuvenation Technologies, Inc., and has equity and stock options in the company. M.T., C.M.M., C.A.H., Y.X.W., K.E.G.M., A.K., and A.C. declare that they have no known competing financial interests or personal relationships that could have appeared to influence the work reported in this paper.

