## Supplementary Information for "Machine learning-based classification of binary dynamic fluorescence signals reveals muscle stem cell fate transitions in response to pro-regenerative niche factors"

1. Supplementary Figures…………………………………………………………………...2

2. Supplementary Video Captions………………………………………………………..4

3. Supplementary Table…………………………………………………………………...5

### Supplementary Figures

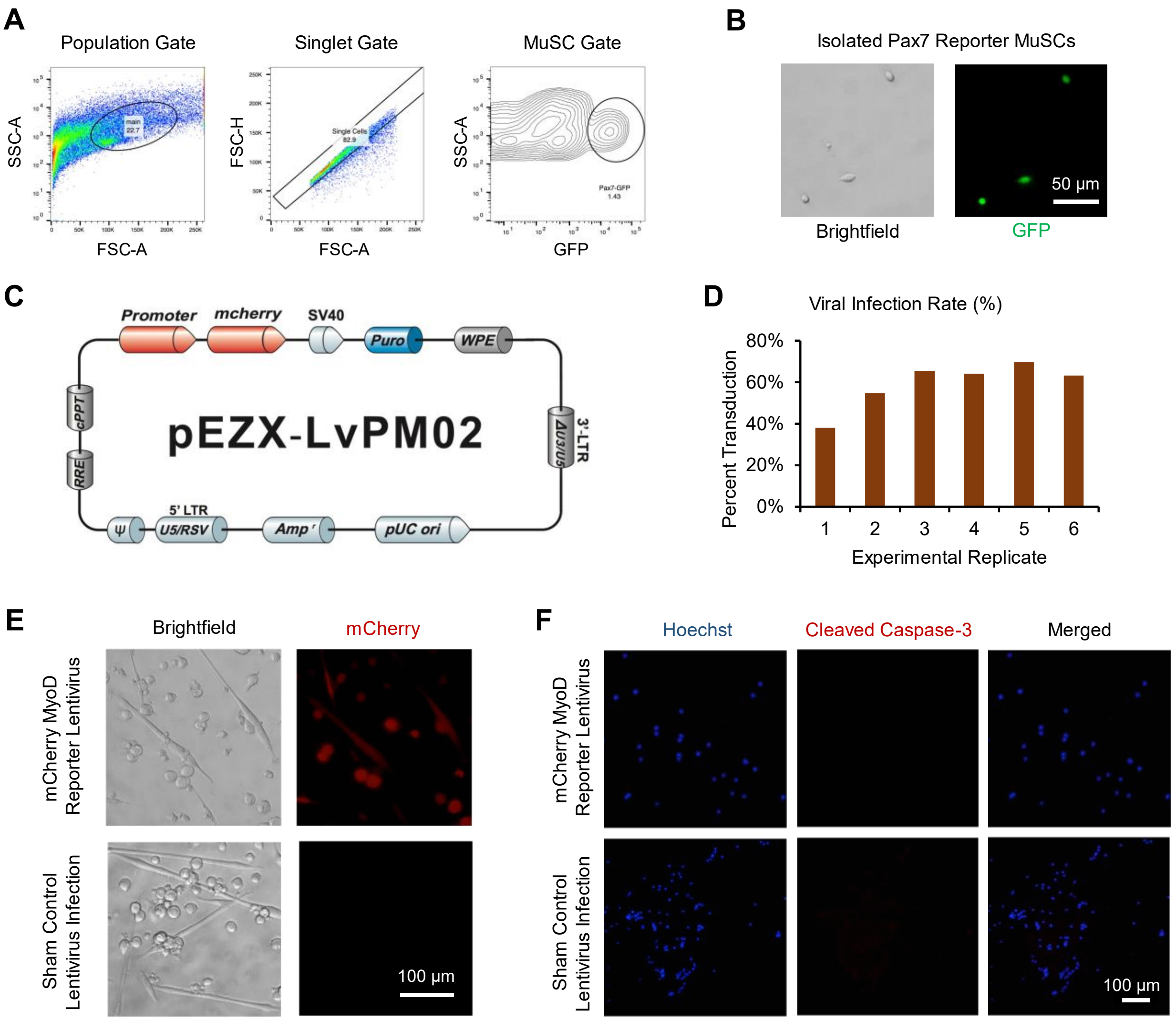


**Figure S1. Characterization of binary fluorescent reporter MuSCs. (A)** Representative FACS gating strategy to enrich for Pax7-eGFP reporter MuSCs. **(B)** Representative brightfield and fluorescence images of sorted Pax7-eGFP reporter MuSCs. **(C)** Plasmid map for the lentiviral vector used to generate the MyoG-mCherry dual reporter MuSCs. **(D)** Viral infection rate across experimental replicates. **(E)** The MyoG-mCherry commitment reporter was validated by inducing differentiation in transduced and sham control MuSCs and measuring mCherry fluorescence by microscopy. **(F)** Transfected MuSCs exhibited negligible viral toxicity as measured by immunofluorescence for the apoptosis marker cleaved caspase-3.

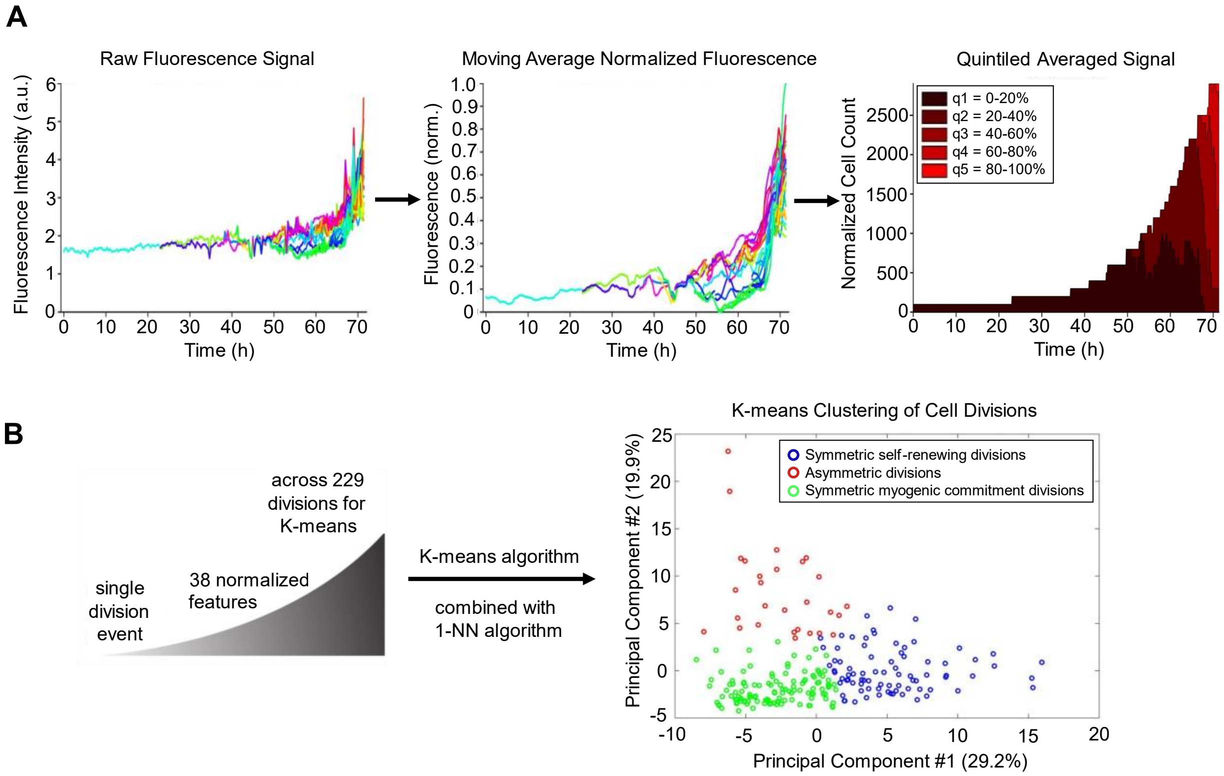


**Figure S2. Automated analysis of fluorescence signals for division classification. (A)** The raw fluorescence intensity signals from a single MuSC and its progeny were normalized to the maximum value in the microwell over the time course experiment and smoothed with a moving average filter. The normalized fluorescence signal was then used to bin the cells into quintiles of reporter expression. **(B)** To classify each division event, 38 normalized features were extracted from the time lapse data (see Supplementary Table S1) and fed into a K-means algorithm to cluster the division features. Cell division assignments were performed using a 1-nearest neighbor (1-NN) classifier.

### Supplementary Video Captions

**Video S1. Representative time lapse video of vehicle treated (control) MuSC colony.** Brightfield frames were taken every 5 minutes, and fluorescence images of the cell fate reporters were taken every 20 minutes. The color of the overlaid fluorescence signal corresponds to the Pax7 GFP reporter (green) or the myogenin mCherry reporter (red).

**Video S2. Representative time lapse video of PGE2 treated MuSC colony.** Brightfield frames were taken every 5 minutes, and fluorescence images of the cell fate reporters were taken every 20 minutes. The color of the overlaid fluorescence signal corresponds to the Pax7 GFP reporter (green) or the myogenin mCherry reporter (red).

**Video S3. Representative time lapse video of OSM treated MuSC colony.** Brightfield frames were taken every 5 minutes, and fluorescence images of the cell fate reporters were taken every 20 minutes. The color of the overlaid fluorescence signal corresponds to the Pax7 GFP reporter (green) or the myogenin mCherry reporter (red).

### Supplementary Table

**Table S1: Extracted features with relative weights for K-means clustering of divisions.** The weights were empirically determined by validation with ground truth lineage trees in which the divisions were manually labeled. The fluorescence quantified signals account for 30 features out of a total of 38.

| **Feature** | **Weight** |
|---|---|
| Parent/daughter myogenin min | 2 |
| Parent/daughter myogenin max | 2 |
| Parent/daughter myogenin avg | 2 |
| Parent/daughter myogenin variance | 2 |
| Parent/daughter myogenin beginning-end span | 2 |
| Parent/daughter Pax7 min | 1 |
| Parent/daughter Pax7 max | 1 |
| Parent/daughter Pax7 avg | 1 |
| Parent/daughter Pax7 variance | 1 |
| Parent/daughter Pax7 beginning-end span | 1 |
| Daughter cells lifetime difference | 3.5 |
| Daughter 1/2 divided? | 1 |
| Daughter 1/2 died? | 1 |
| Daughter cells fate difference (division) | 1 |
| Daughter cells fate difference (death) | 1 |
| Parent generation | 1.5 |
